# *Pseudomonas* isolates degrade and form biofilms on polyethylene terephthalate (PET) plastic

**DOI:** 10.1101/647321

**Authors:** Morgan Vague, Gayle Chan, Cameron Roberts, Natasja A. Swartz, Jay L. Mellies

## Abstract

Bioaugmentation is a possible remediation strategy for the massive amounts of plastic waste in our oceans and landfills. For this study, soil samples were collected from petroleum polluted locations in the Houston, Texas area to isolate microorganisms capable of plastic degradation. Bacteria were propagated and screened for lipase activity, which has been associated with the bacterial degradation of some plastics to date. We identified three lipase-positive *Pseudomonas* species, and *Bacillus cereus* as part of two consortia, which we predict enhances biofilm formation and plastic degradation. Lipase-positive consortia bacteria were incubated alongside blank and *E.coli* controls with UV-irradiated polyethylene terephthalate (PET), high-density polyethylene (HDPE), or low-density polyethylene (LDPE) as sole sources of carbon. Surface degradation of PET plastic was quantified by changes in molecular vibrations by infrared spectroscopy. The bacteria formed biofilms on PET, observed by scanning electron microscopy, and induced molecular changes on the plastic surface, indicating the initial stages of plastic degradation. We also found molecular evidence that one of the *Pseudomonas* isolates degrades LDPE. To date, lipase positive *Pseudomonas* spp. degradation of PET has not been well described, and this work highlights the potential for using consortia of common soil bacteria to degrade plastic waste.

## INTRODUCTION

It is estimated that 300 million tons of plastic waste is generated every year, with 30-33 million tons originating in the US alone ^1^. This number, however, underestimates the plastic burden on the planet as it does not reflect the millions of tons of waste that go unreported each year ^2^. The plastics industry is projected to continue its growth, with profits expected to exceed $375 billion annually by 2020 as plastic begins to overtake the medical device sector, and singleuse food and beverage packaging continues to dominate the international food landscape ^3^. This is worrisome for many reasons. Over 50% of plastic produced internationally in 2014 went toward single-use plastic food and beverage packaging, which was quickly discarded as waste rather than recycled ^3^. This plastic waste accumulates in landfills and oceans, where it can persist for many decades, though estimates for persistence vary. Of the 8.3 billion metric tons of plastic that have been produced since their introduction to the consumer market following WWII, roughly 6.3 billion metric tons are estimated to have become plastic waste, with 79% accumulating in landfills, and 19% ending up in our oceans ^1,4^. This number is most likely an underrepresentation of the plastic currently residing in the oceans as environmental researchers have recently determined that the majority of plastic debris in the ocean resides in deep sea sediments, which act as a plastic sink ^5^.

Polyethylene terephthalate (PET) plastic, the focus of this work, is composed primarily of repeating ethylene glycol and terephthalic acid (TPH) monomers. PET’s linear structure and high proportion of aromatic components are chemically inert and increase its durability, making it highly resistant to degradation ^6^. Its rigid structure and ability to form an effective gas barrier against molecular oxygen make PET a popular choice for use in water bottles and single-serving containers. It is the most commonly produced polymer worldwide, and while its presence in water bottles has been popularized, it is also used in common household goods such as carpet fibers, curtains and fabrics. Polyethylene (PE) and derivatives are a group of polymers that are relatively hydrophobic, chemically inert, possess a high molecular weight and sometimes a branched 3D structure. Each of these shared characteristics reduce their likelihood of being used as a carbon source for microorganisms ^7^. Largely resistant to biodegradation, PE and its derivatives can persist in the environment for multiple decades, depending on polymer type ^8^.

Biodegradation is the process by which microorganisms, usually bacteria or fungi, induce polymer degradation via assimilation or the release of enzymes that can cleave various molecular bonds within the polymer. Spontaneous hydrolysis, photo-oxidation, and mechanical separation of plastic have been shown to enhance biodegradation by introducing cleavable bonds or simply increasing plastic surface area for colonization ^9^. Generally, any microorganism capable of reducing plastic polymers to CO_2_ and water (aerobic conditions), CO_2_ and methane (anaerobic conditions), or inorganic molecules and biomass is considered capable of biodegradation. Biodegradation occurs through four types of processes: solubilization, ionization, hydrolysis and enzymatic cleavage. Solubilization is essential for biodegradation because increased solubility broadens the availability of polymers to biodegrade organisms. Enhancing biodegradation with the use of synthetic or biosurfactants, which can increase the solubility of polymers, have been suggested as a component of bioremediation strategies ^10,11^. Ionization is solubilization brought on by protonation or ionization of a side-chain group present on certain polymers and induced by changes in pH. Spontaneous hydrolysis of side-chain ester bonds can lead to solubilization, while hydrolysis of polymer backbone ester bonds leads to degradation. For hydrolysis to occur, the polymer must contain ester bonds that can be attacked by water and cleaved. For instance, polyester, which contains multiple ester linkages in each monomer, is degraded primarily by spontaneous hydrolysis ^12^. Accordingly, synthetic polymers such as PE, polypropylene (PP), and polystyrene (PS) lack natural ester functionality and are resistant to hydrolysis unless they first undergo oxidation.

Enzymatic cleavage of plastic polymers by microbial or fungal enzymes requires two steps. Initially, secretion and adhesion of the enzyme to the polymer surface is followed by catalysis of primarily ester bond cleavage. Due to this mechanism, plastics that are susceptible to microbial enzyme degradation must either naturally contain ester bonds or be oxidized by another method prior to catalysis. Accordingly, pre-treatment by UV radiation to induce ester functionality via photo-oxidation greatly increased the ability of *Brevibacillus borstelensis* to degrade polyethylene, and for low-density polyethylene (LDPE) films to be degraded by a mixed culture of *Lysinibacillus xylanilyticus* and *Aspergillus niger* ^7,13^. In addition, two *Penicillum* spp. have been shown to have HDPE and LDPE degrading capabilities ^14^. PET on the other hand, is more-readily degraded due to its inherent ester linkages by PETase, an enzyme found in the bacterium *Iadonella sakaiensis* and cutinases found in multiple fungal species ^6^.

Once large complex polymers are unzipped into monomers, oligomers, aldehydes, ketones and other small molecules, they can be absorbed into the cell and used as a source of carbon and energy. Mineralization occurs when these degradation products are reduced to the end products of carbon dioxide, methane, and water. Released CO_2_ is then absorbed by plants or photosynthetic bacteria. The process of photosynthesis and carbon fixation returns carbon from plastic to the biosphere.

We predict that if plastic degrading bacteria are more ubiquitous than previously thought and can be readily isolated from petroleum-contaminated sites, then these bacteria can be utilized for a bioaugmentation strategy to reduce plastic waste. While it is known that certain bacteria can degrade plastics, PET degradation by lipase positive *Pseudomomas* spp. has not been described well ^15^. Further, little has been done to investigate how bacterial consortia might be utilized for bioaugmentation to optimize degradation and to mitigate PET, or other plastic pollution.

## RESULTS

### Bacterial Isolation and Screening

To screen isolates for lipase activity, cultures on master plates of LB agar were colony-stamped directly onto Rhodamine B agar plates, incubated and subjected to 365 nm UV light (Fig. 1). For Rhodamine B plates, olive oil was used as a source of long-chain fatty acids to check for lipase activity. Rhodamine B dye binds to free fatty acids (cleaved by a secreted lipase) and glows when exposed to UV radiation (Fig. 1D). Thus, the presence of glowing halos around the colonies is indicative of lipase activity. Colonies that appeared lipase-positive by stamping were isolated and re-tested to confirm lipase activity and to isolate pure cultures. In total, 192 colonies were screened after appearing lipase-positive (or growing near lipase-positive bacteria) during initial Rhodamine B screening.

**Figure 1.**
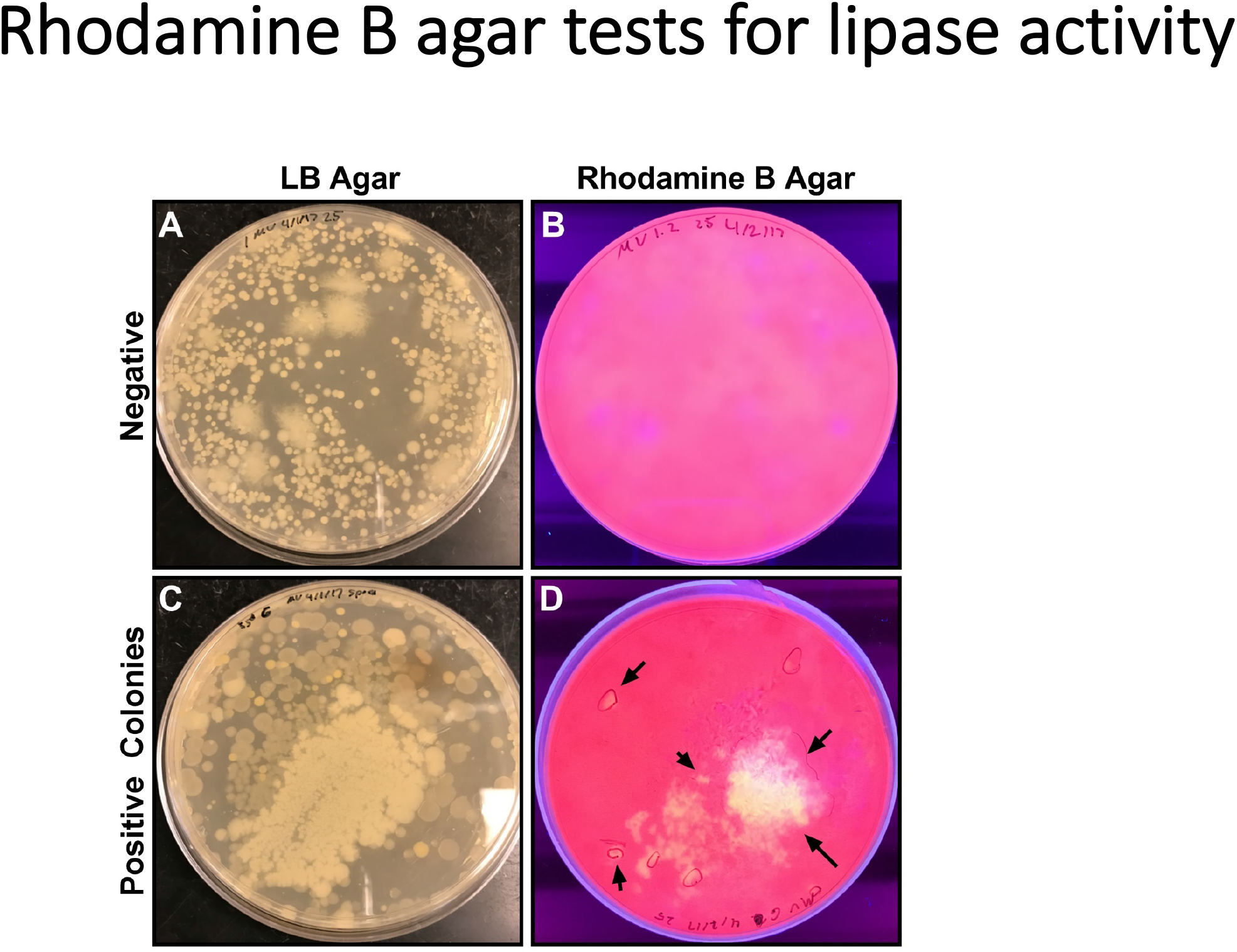
Rhodamine B Agar screen for lipase activity. Master plates of mixed colonies were generated by soaking soil samples in water and collecting the supernatant to spread on LB plates. Individual plates with growth (representative plates in A and C) were stamped onto Rhodamine B plates (4% w/v) (B and D) to screen for lipase activity. The presence of orange or yellow halos under 365nm UV exposure indicates lipase positive colonies (indicated with arrows). After, individual colonies in lipase positive areas were spotted onto new Rhodamine plates to isolate the lipase producers, and positive spots were re-streaked onto LB agar for purification.

Each colony was repeatedly streaked for isolation and tested via serial Gram staining for purity until pure isolates were obtained (Fig. 2). Initial Gram stains indicated the cultures were mixed, with varying morphologies, most likely as consortia on the LB agar plates. Therefore, multiple re-streaking was performed. From Consortium 9, two isolates were purified, a Gram-negative rod (Isolate 9.2) and a Gram-positive rod (Isolate 9.1) (Fig. 2). From Consortium 13, two isolates were also purified, a Gram-negative rod (Isolate 13.2) and a Gram-positive rod (Isolate 13.1). Consortium 10 eventually had a single morphology by Gram stain indicating the consortium was purified to a single isolate (Isolate 10). The Rhodamine B screen using master plates allowed for screening of thousands of colonies for lipase activity, but positive cultures were mixed and thus subsequently reduced to single isolates by streaking to pure culture, tracking progress using Gram staining and colony morphology.

**Figure 2.**
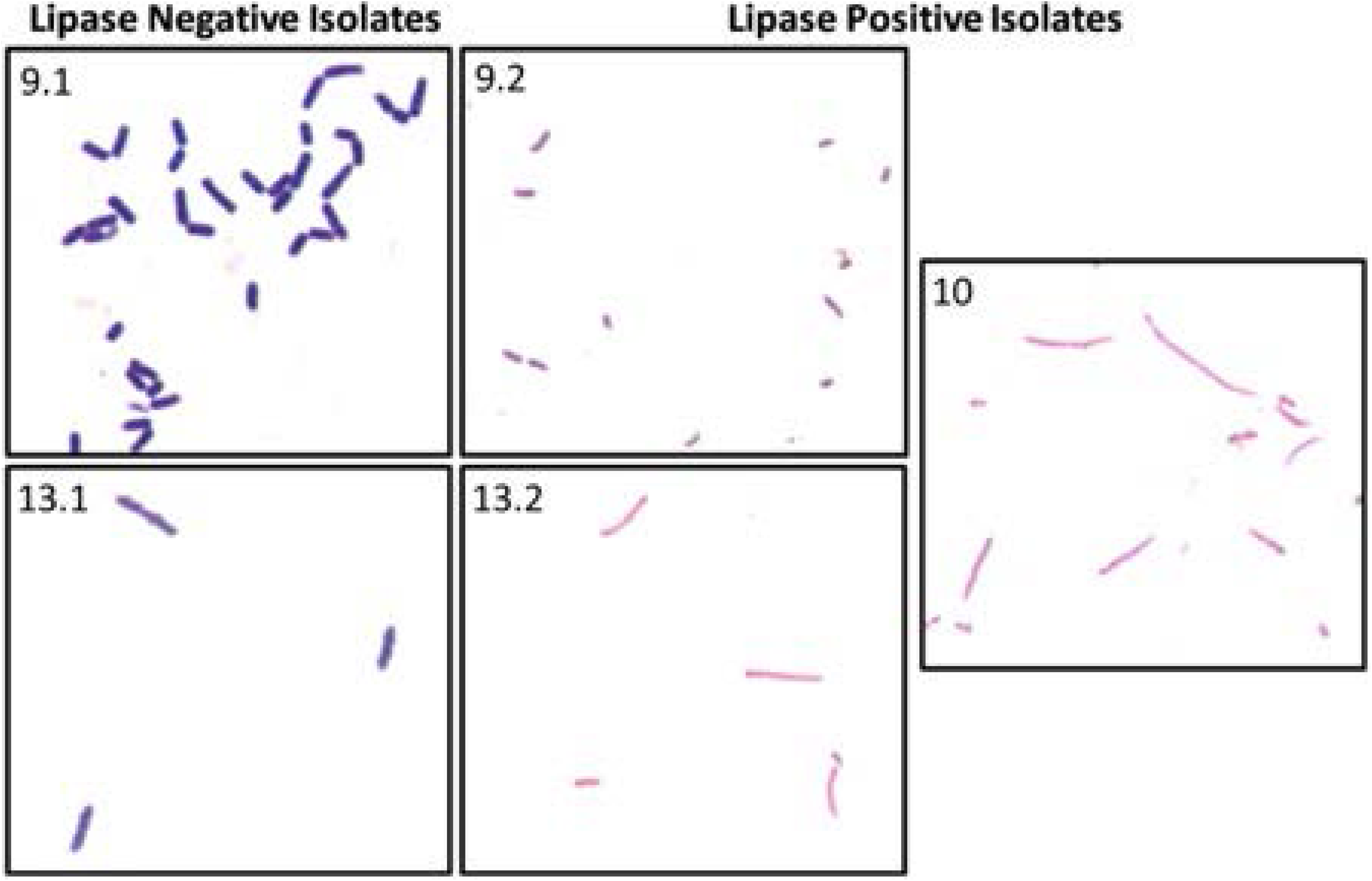
Gram stains of pure isolates from lipase-positive consortia. Lipase negative isolates were Gram positive rods (left) while all three lipase positive isolates were Gram negative rods (right). Isolates 10 and 13.2 were rods that were more elongated than Isolate 9.2. Scale bar = 10 μM.

Pure cultures were tested for lipase activity using Rhodamine B agar plates (Fig. 3). Isolates 9.2, 10 and 13.2, all Gram-negative rods, tested positive for lipase activity, suggesting that they might be capable of plastic degradation (Figs. 3D). Isolates 9.1 and 13.1 were lipase negative, as was the negative control, *E. coli* strain MC4100. Fluorescent halos, indicating lipase activity, on Rhodamine B agar can be easily distinguished from colony growth and any natural fluorescence produced by Pseudomonads (Figs. 3A and C). Because of the halo consistently observed for Isolate 9.2 to be larger than that of the other isolates under UV light at 365 nm (Fig. 3D), the data suggested greater lipase activity for this isolate compared to the other two lipase producers, Isolates 10 and 13.2. These results confirm that the Rhodamine B agar test can be used to identify lipase positive, and putative plastic degrading bacteria.

**Figure 3.**
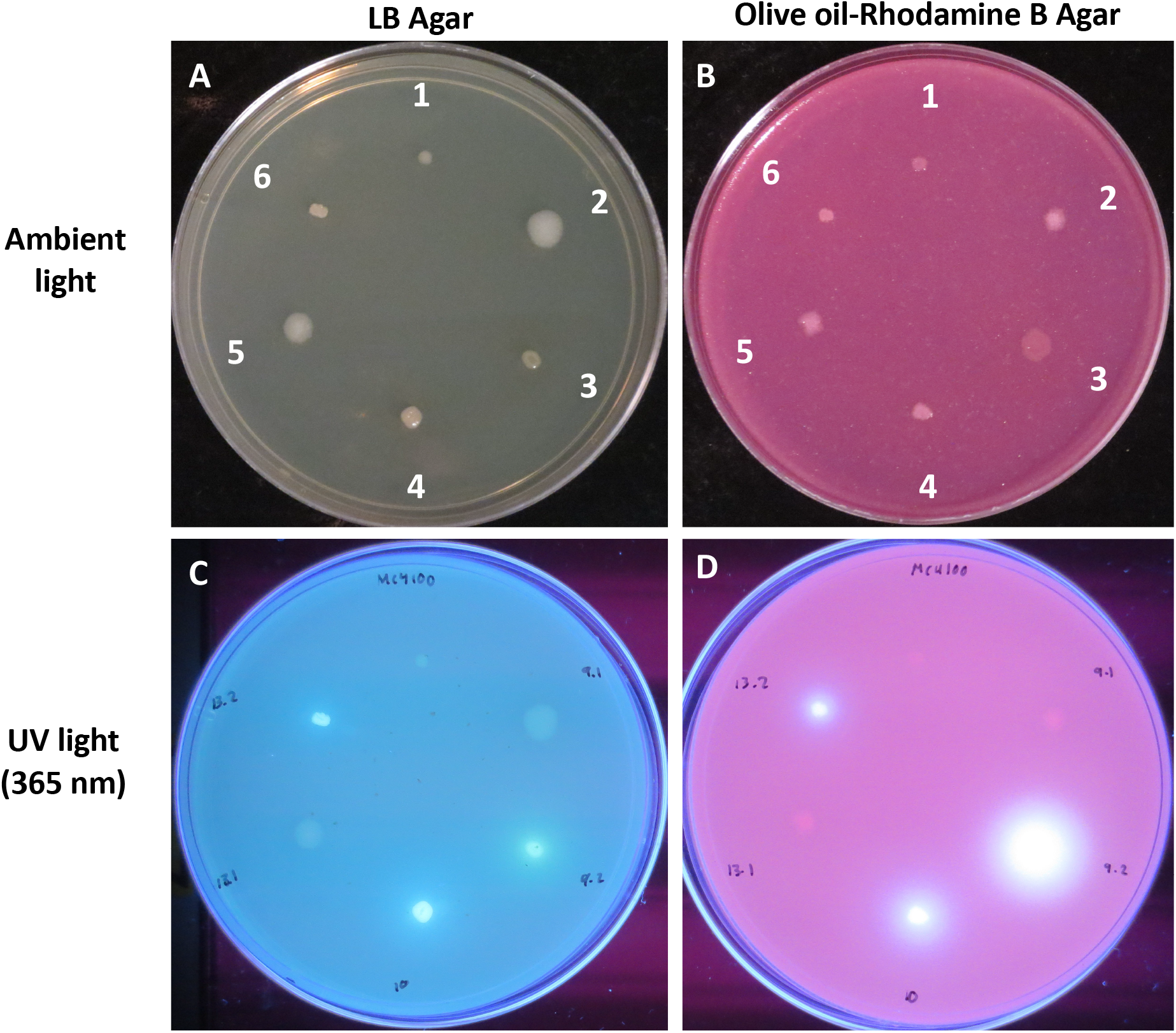
Lipase production of isolates plated on Rhodamine B agar containing olive oil (See Materials and Methods) demonstrating lipolytic activity. Lipase negative *E coli* strain MC4100 (1) was a negative control. Isolates 9.1 (2), 9.2 (3), 10 (4), 13.1 (5), and 13.2 (6) were inoculated on LB and olive oil-Rhodamine B plates and allowed to grow for 48hr at 26°C. The LB and Rhodamine B plates with no UV exposure are shown in A and B. LB and Rhodamine B plates exposed to UV light at 365 nm are shown in C and D. The LB plates show natural fluorescence (C) and the Rhodamine B plate (D) shows lipolytic activity indicated by white halos for isolates 9.2, 10, and 13.2. Images in A and C are the same plate photographed without and with UV light, respectively. Images in B and D are the same plate photographed without and with UV light, respectively. The original uncropped images (**Fig. S2**) are provided in the **Supplementary Information**.

### Identification of lipase positive and consortia isolates

Taxonomic dentification of pure isolates was done by 16S rRNA gene sequencing. All three lipase positive isolates were identified as *Pseudomonas* with 100% identity (E value = 0.0). Pseudomonads often produce fluorescent compounds (see Figure 3C), and thus this was consistent with the 16S rRNA gene sequencing data. The consortia members lacking lipase activity were *Bacillus* spp. with 100% identity (E value = 0.0). 16S rRNA genes from Isolate 9.2 and Isolate 10 were amplified from pure isolates, while that from 13.2 had been previously identified from Consortium 13. A refined analysis was necessary to identify the bacteria at the species level. Thus, amplification of the variable regions, V3 to V6 or the rRNA genes, was employed ^17^. Isolates 10 and 13.2 were determined to be *Pseudomonas putida* (99% identity, E values = 0.0). Isolate 9.2 was determined to be *Pseudomonas chlororaphis* (99% identity, E value = 0.0). Consortia members of 9.1 and 13.1 were identified as *Bacillus cereus* using this method (99% and 100% identity, respectively, E values = 0.0).

To determine whether the *Pseudomonas putida* and *Bacillus cereus* species were unique isolates, we employed RAPD. As seen in Figure 4B, isolates 9.2, 10, and 13.2 exhibited unique RAPD patterns. Similarly, the non-lipase containing 9.1 and 13.1 were unique isolates of *Bacillus cereus.* Thus, the Consortium 9 members were *Pseudomonas chlororaphis* and *Bacillus cereus* subspecies 9.1; Isolate 10 was *Pseudomonas putida*; and Consortium 13 contained *Pseudomonas putida* subspecies 13.2 and *Bacillus cereus* subspecies 13.1

**Figure 4.**
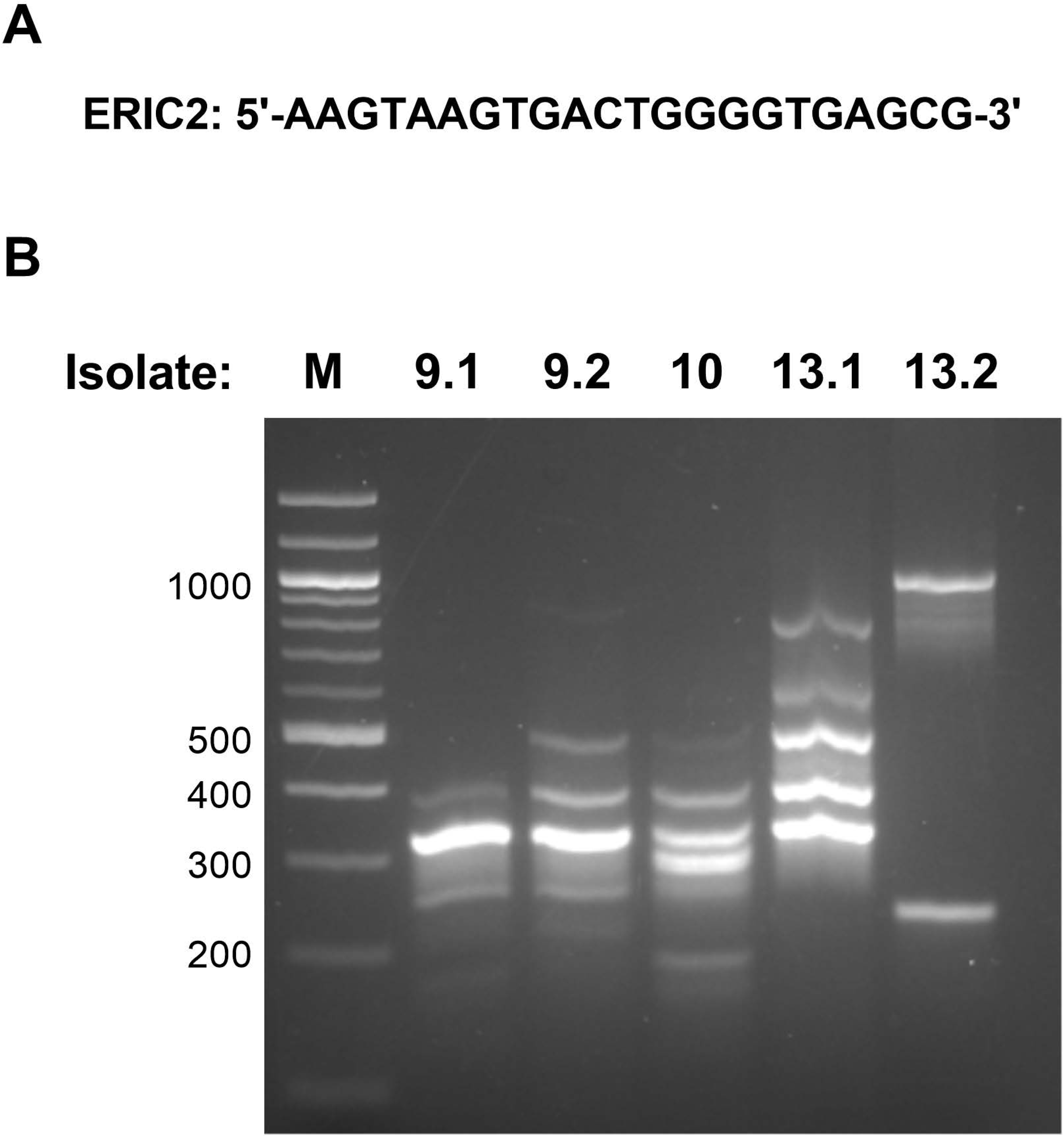
RAPD PCR analysis. RAPD PCR was conducted on each of the five bacterial isolates using the A) ERIC2 primer (5’-AAGTAAGTGACTGGGGTGAGCG-3’), with 57° C annealing temperature^17^. B) PCR products were imaged in a 1.8% agarose gel. All isolates were determined to be unique by this analysis.

### Plastic degradation

While lipases are commonly identified plastic-degrading enzymes, the presence of a lipase is suggestive, not conclusive of whether an isolate is capable of degrading plastic ^6^. Liquid cultures were set up in triplicate in carbon free media inoculated with each lipase positive consortium (Consortia 9 and 13, and Isolate 10) or *E.coli* MC4100 as a negative control. Sterilized strips of PET, HDPE, or LDPE were placed in each tube to be used as a carbon source, with all other nutritional requirements provided in the media. Half of the samples were pretreated with UV radiation, which has been shown to enhance plastic degradation via the introduction of lipase-cleavable ester bonds ^7,13^.

Given that degradation was likely to occur on the surface as opposed to the bulk material, a surface-sensitive technique such as ATR-FTIR is ideally suited to monitor chemical changes of degraded plastics ^18,19^. The averaged spectrum collected for UV-irradiated PET incubated with positively isolated bacteria (13uv) and without (Buv) are shown in Figure 5A where major vibrational modes and respective chemical bonds of the polymer identified. Difference spectra for PET (Fig. 5B) were calculated by subtraction of the averaged method blank spectrum from each averaged treatment method spectrum, where arrows indicate growth (↑) or loss (↓) of chemical functionality in the polymer. As plastic degrades, additional ester (C=O, C-O), carboxyl (C=O, C-O, O-H), alcohol (C-O, O-H), and terminal vinyl (=CH_2_) groups are created in the polymer where the appearance or alteration of these bonds will change peak intensities at 2958 cm^−1^, 1713 cm^−1^, 1089 cm^−1^, 888 cm^−1^, and 730-710 cm^−1^. Changes in polymeric bonds indicating degradation can be quantified by using an index calculated from the ratio of characteristic peak intensities to the normal C-H bending mode (at 1409 cm^−1^in PET). For instance, carboxylation of shortened hydrocarbon chains by photo-oxidation occurs prior to the β-oxidation cycle; and therefore, a low carbonyl index (decrease in ester C=O) combined with new C=O and O-H peaks suggests bacteria were actively converting plastic into precursors for β-oxidation or the TCA cycle. Using peak intensities, the relative amounts of aliphatic, carbonyl, and ester functionality of the plastic are listed in Table 1 for comparison of PET samples with and without inoculation or pretreatment by UV irradiation. Only the UV pretreated PET samples showed substantial, reproducible spectral changes when analyzed by ATR-FTIR.

**Figure 5.**
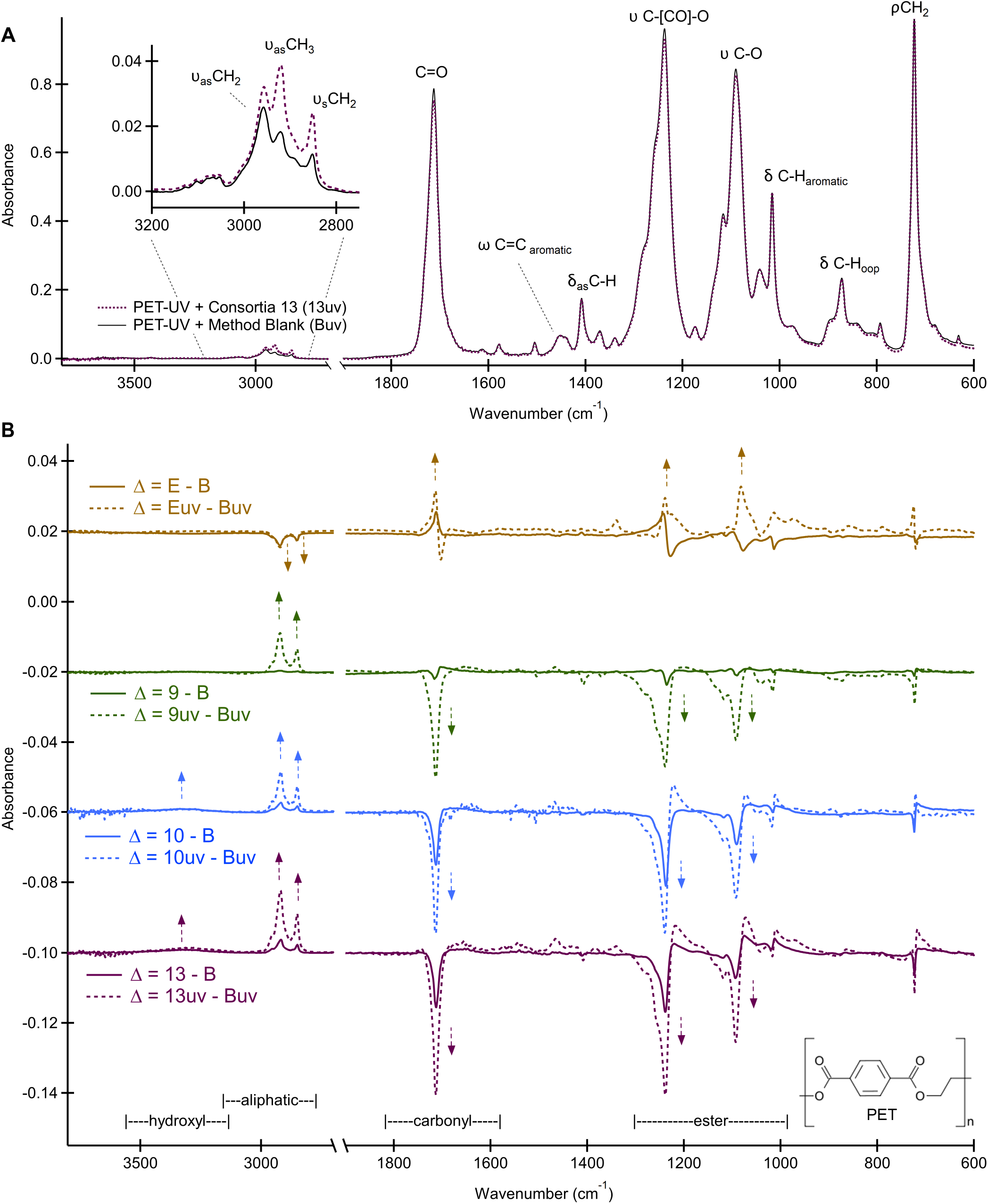
Infrared spectra of PET plastics incubated with lipase positive consortia 9 and 13, isolate 10, and *E,coli.* A) Averaged ATR-FTIR spectrum acquired of the method blank (Buv) as compared with consortium 13 (13uv). B) Comparison of PET difference spectra with UV (dashed line) and without UV (solid line) pretreatment prior to inoculation. Difference spectra (Δ) are vertically offset for clarity and were produced by spectral subtraction of the method blank (Buv) from the inoculated PET (9uv, 10uv, 13uv, or Euv). Direction of arrows indicate the growth or loss of a peak after incubation, signifying a relative increase or decrease in abundance of that bond, respectively. All spectra were averaged (n = 9) and normalized by PA to δ_as_ at 1408 cm^−1^ prior to spectral math.

**Table 1.**
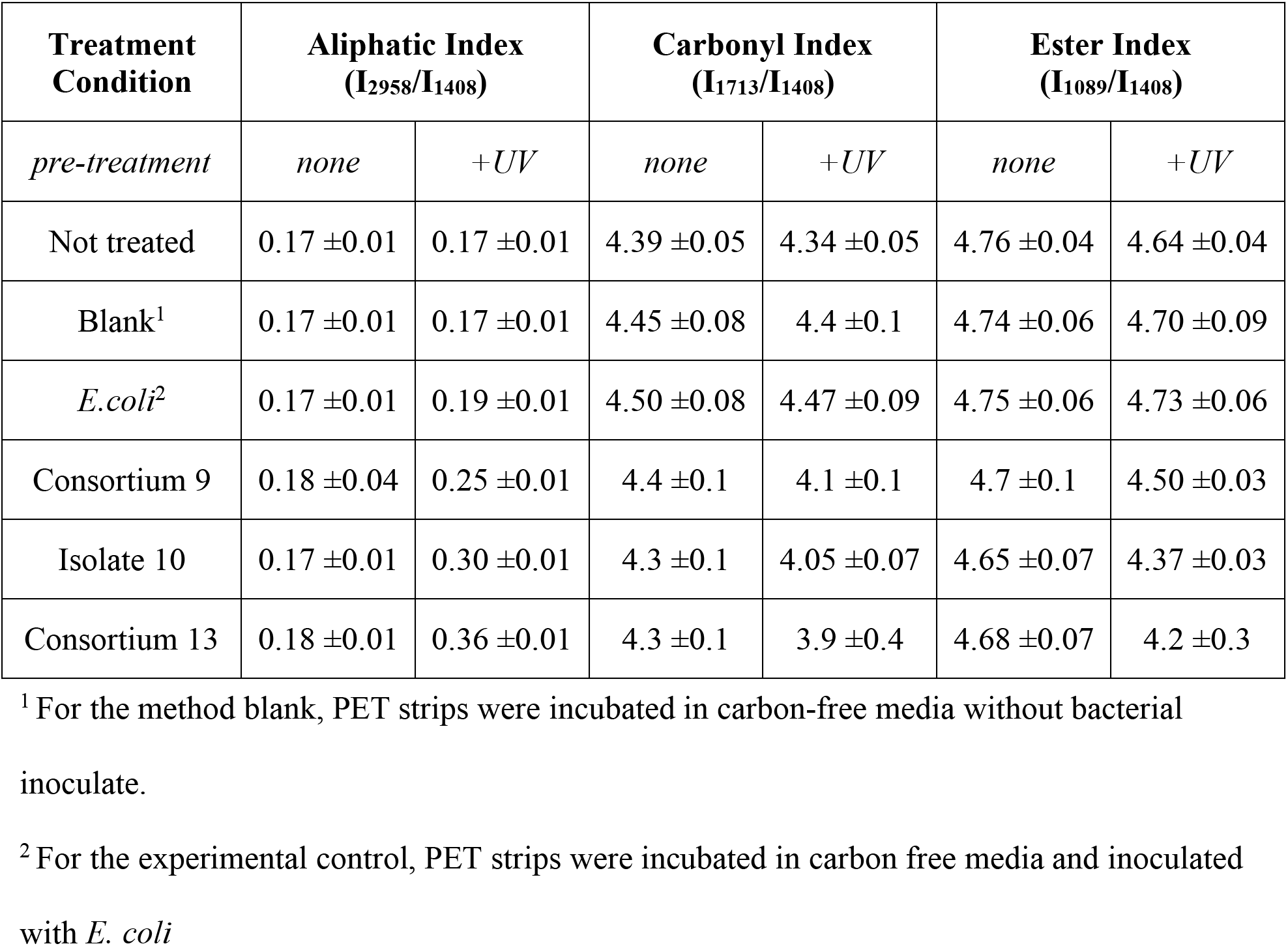
Molecular index and confidence intervals calculated by relative intensities of vibrational bands from ATR-FTIR analysis of commercial PET with and without UV-pretreatment incubated in carbon-free media for six weeks (n = 9, α = 0.05).

Decreases in the carbonyl and ester indexes are expected during PET degradation as C=O and C-O bonds are broken during ester-cleavage. If undergoing chain scission of the polymer network, a simultaneous increase in the aliphatic index is expected as the abundance of saturated terminal ethylene glycol groups increases, and mid-chain methylene bonds decrease. These changes indicating degradation of the plastic surface were observed for PET incubated with all three positively identified lipase-producing strains, as shown in Table 1. Typical markers of plastic degradation by the lipase-positive bacteria were observed where the carbonyl index decreased systematically (when compared to Buv = 4.4 ±0.1 or Euv = 4.47 ±0.07, 9uv = 4.1 ±0.1, 10uv = 4.05 ±0.07, and 13uv = 3.9 ±0.4), as did the ester index (when compared to Buv = 4.70 ±0.09 or Euv = 4.73 ±0.06, 9uv = 4.50 ±0.03, 10uv = 4.37 ±0.03, and 13uv = 4.2 ±0.3). Suspected ester-cleavage and chain-scission occurring on the surface of these samples can be visualized in the difference spectra (Fig. 5B). Like the directionality of the calculated indexes, in general, a positive peak denotes a new product and negative peak denotes loss. Surprisingly the aliphatic index appeared most sensitive to treatment method and changes can be clearly identified in the difference spectra from 3200-2800 cm^−1^. All lipase-positive inoculated PET samples presented a substantial increase in their aliphatic index (9uv = 0.25 ±0.01, 10uv = 0.30 ±0.01, 13uv = 0.36 ±0.01) when compared to control samples (Buv = 0.17 ±0.01 and Euv = 0.19 ±0.01). Loss of ester functionality was confirmed by the negative peaks in all lipase-positive difference spectra of PET (Fig. 5B), whereas the *E.coli* samples showed a systematic increase or enrichment in the same region between 1300 – 1000 cm ^−1^. Even though small changes were observed in the averaged difference spectrum for Euv samples, no significant difference was found between carbonyl and ester indexes of blank and *E. coli* incubated PET samples (Tukey HSD: Euv to Buv, p_carbonyl_ = 0.946 and p_ester_ = 0.997). The aliphatic index of *E.coli* samples was observed to be significantly smaller than blank samples (Tukey HSD: Euv to Buv, p_aliphatic_ = 0.0356). In contrast, lipase-positive PET samples 9uv, 10uv, and 13uv were found to have substantially larger aliphatic indexes accompanied by a simultaneous decrease in carbonyl and ester indexes that suggest decarbonylation was occurring. These data indicate spontaneous hydrolysis of PET in the carbon-free media was not a favored mechanism during incubation and any observed changes were instead due to the bacteria selected for inoculation. Overall, the decrease in the carbonyl and ester indices typically used as markers for plastic degradation were greatest in the UV pretreated PET samples inoculated with lipase-positive bacteria, suggesting UV treatment and microorganism biodegradation are synergistic for PET plastics.

Difference spectra of LDPE samples under the same treatment conditions as PET are included in supplementary information (Fig. S1) along with calculated molecular indices for both LDPE and HDPE (Table S1 and S2, respectively). Only Isolate 10 showed similarly promising results in degrading both PET and LDPE as substantial changes were not observed in any lipase-positive HDPE samples. Most notable in the averaged spectrum of LDPE incubated with Isolate 10 as compared to the blank (Fig. S1A) is an increase in vibrational modes associated with O-H stretching and bending at 3400 and 985 cm^−1^, respectively. The addition of O-H groups may be related to the addition of primary or secondary alcohols on the polymer backbone, an early step identified in the biodegradation of PE ^20^. The addition of alcohol or ethylene glycol groups in LDPE after incubation with Isolate 10 was confirmed by an increase in new C-O vibrational modes from 1300-1000 cm^−1^ (Fig. S1B). The average aliphatic index of LDPE samples incubated with Isolate 10 also increased when compared to blank and *E.coli* samples, but only for samples that were not pretreated with UV radiation (10 = 31 ±1, E = 26 ±2, B = 27±2). The increase in aliphatic functionality in Isolate 10 samples was accompanied by a decrease in carbonyl index (10 = 0.33 ±0.05, E = 0.38 ±0.05, B = 0.37 ±0.05), and increase in methylene index (10 = 15 ±1, E = 13 ±1, B = 13 ±1). In contrast to PET, these data indicate that for plastics composed primarily of LDPE, UV treatment and microorganism biodegradation are not necessarily synergistic.

Our data also suggested that the consortia could have varying capabilities to colonize and degrade the plastic or to produce lipase compared to the individual *Pseudomonas* isolates. To test this idea, we cultured Consortium 9 and individual isolates, taken from bacteria cultured on PET, then inoculated onto Rhodamine B plates to quantify lipase activity. Measuring the ratio of fluorescent halo to growth^16^, Consortium 9 possessed greater lipase activity than either Isolate 9.2 or 10 alone (p < 0.001; Table 2). These data indicated that *P. chlororaphis* cultivated with *Bacillus cereus* produced and/or secreted greater quantities of lipase compared to *P. chororaphis* or *P. putida* alone.

**Table 2.**
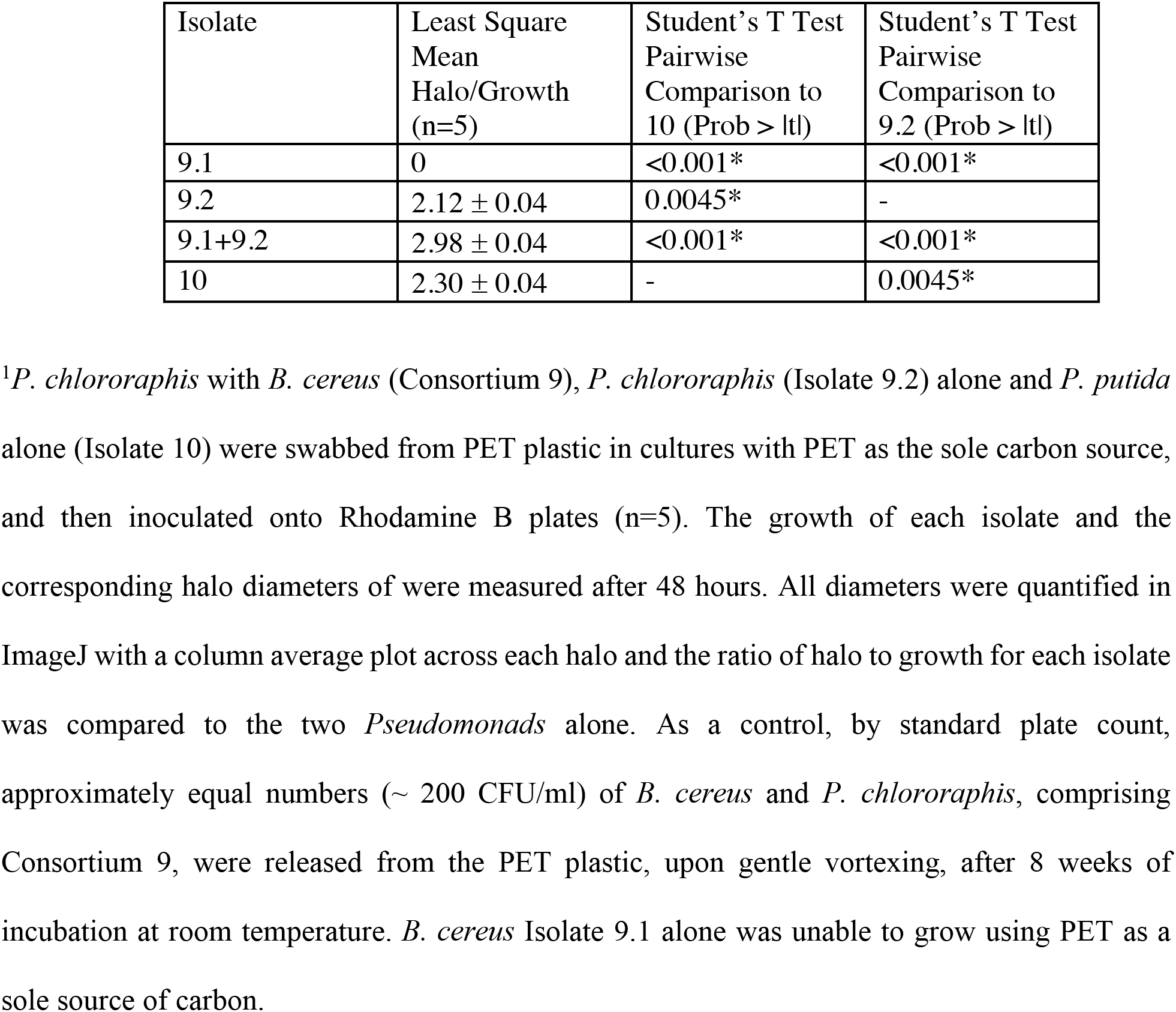
Ratios of lipase production to colony growth for Consortium 9 and individual isolates^1^.

| Isolate | Least Square Mean Halo/Growth (n=5) | Student's T Test Pairwise Comparison to 10 (Prob > \|t\|) | Student's T Test Pairwise Comparison to 9.2 (Prob > \|t\|) |
| --- | --- | --- | --- |
| 9.1 | 0 | <0.001* | <0.001* |
| 9.2 | 2.12 ± 0.04 | 0.0045* | - |
| 9.1+9.2 | 2.98 ± 0.04 | <0.001* | <0.001* |
| 10 | 2.30 ± 0.04 | - | 0.0045* |

<sup>1</sup>*P. chlororaphis* with *B. cereus* (Consortium 9), *P. chlororaphis* (Isolate 9.2) alone and *P. putida* alone (Isolate 10) were swabbed from PET plastic in cultures with PET as the sole carbon source, and then inoculated onto Rhodamine B plates (n=5). The growth of each isolate and the corresponding halo diameters of were measured after 48 hours. All diameters were quantified in ImageJ with a column average plot across each halo and the ratio of halo to growth for each isolate was compared to the two *Pseudomonads* alone. As a control, by standard plate count, approximately equal numbers (~ 200 CFU/ml) of *B. cereus* and *P. chlororaphis*, comprising Consortium 9, were released from the PET plastic, upon gentle vortexing, after 8 weeks of incubation at room temperature. *B. cereus* Isolate 9.1 alone was unable to grow using PET as a sole source of carbon.

### Evidence of biofilm formation

Biofilm formation is essential for colonization of the plastic by microorganisms and without them, plastic cannot be effectively degraded. SEM allows for the visualization of bacterial colonization and biofilm architecture including extracellular polymeric substance (EPS) deposits which are essential scaffolding for productive biofilms ^21^. Both lipase-positive consortia and Isolate 10 were able to colonize and form biofilms on PET, to different extents (Fig. 6). Consortium 13 had fewer adherent cells and less EPS deposits on the PET, indicating a reduced ability to form a biofilm on the plastic (Figs. 6A and D). Interestingly, Consortium 13 was the only consortium without evidence of pili, suggesting 1) that these pili are essential for robust biofilm formation and 2) that this *Pseudomonas putida* isolate might lack or have mutation(s) in the genes necessary to form pili, preventing robust biofilm formation (Fig. 6A). Pili permit attachment to the plastic, and adherence between adjacent cells (yellow and red arrows in Fig. 6C, respectively), facilitating colony formation on hydrophobic plastic surfaces. The biofilm characteristics for each consortium are summarized in Figure 6D, where the blank sample lacked the same indicators of biofilm formation found on bacteria samples. These results demonstrate that, using a lipase screen, biofilm-forming, plastic degrading bacteria can be isolated from petroleum-contaminated soil samples.

**Figure 6.**
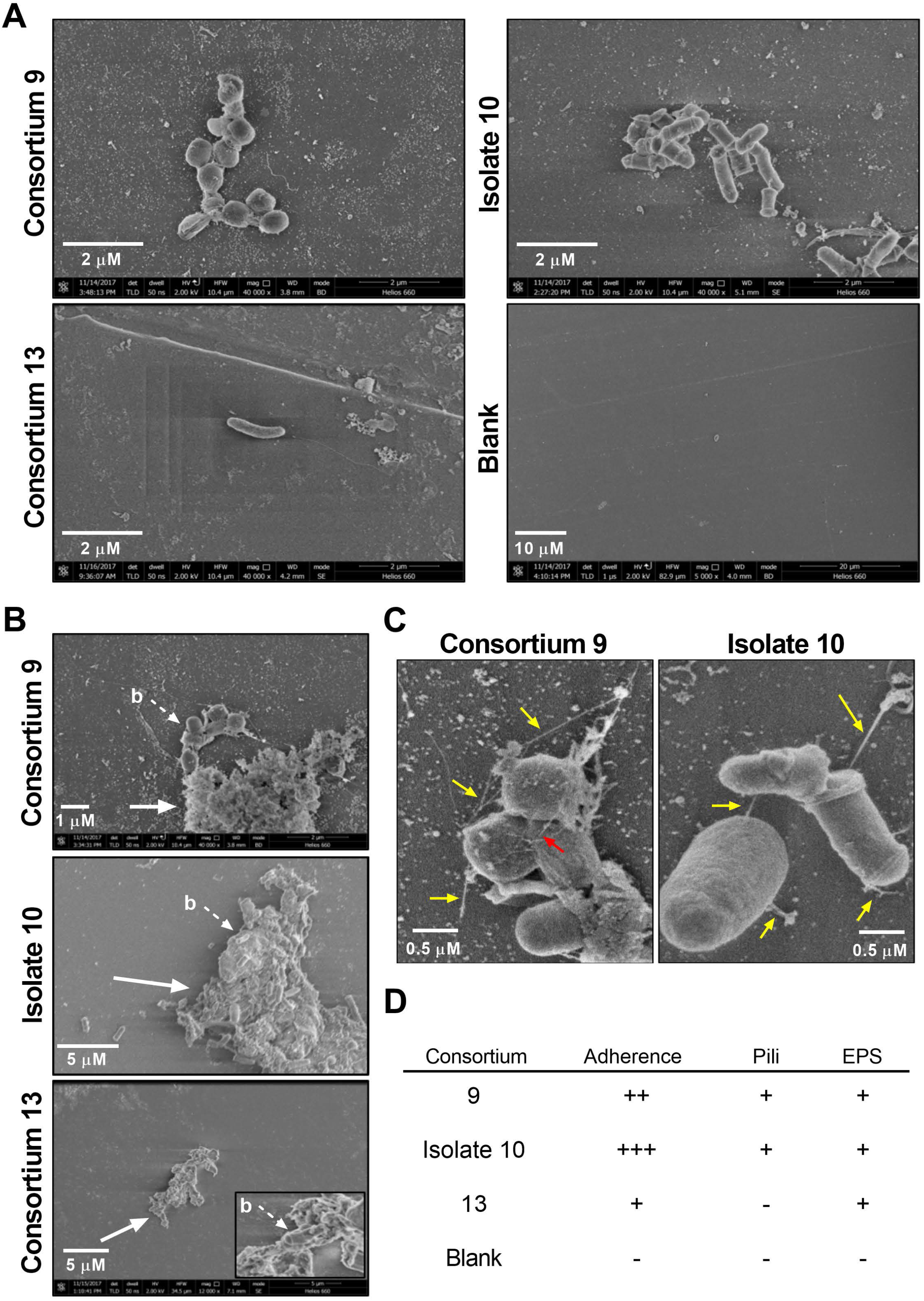
SEM images indicating biofilm formation on PET strips incubated in carbon-free media inoculated with lipase positive consortia and Isolate 10. A) Colonization of PET plastic by Consortium 9, Isolate 10 and Consortium 13 compared with the method blank PET without treatment (blank), which showed no adherence or other hallmarks of bacterial colonization. B) Extracellular polymeric substance (EPS) deposits (white arrow) secreted as part of biofilm formation. Individual bacteria (b, dashed arrow) can be seen embedded in the EPS, which gives biofilms structural integrity. C) Pili formation by Consortium 9 and Isolate 10, aiding in bacterial adherence and biofilm formation. Yellow arrows denote pili attached to PET plastic while red arrow denotes a pilus between bacteria, aiding in cell-cell adhesion. D) Summary of the biofilm morphology observed in multiple SEM images of each consortium. For pili and EPS, (+) denotes presence and (-) denotes absence of given structure. For adherence, consortia were graded from excellent adherence (+++) to poor but observable adherence (+).

### DISCUSSION

Due to their highly stable polymeric structure, plastics do not degrade easily, and strategies must be implemented to assist and enhance their degradation. Accordingly, some in the scientific community have focused on plastic-degrading microorganisms as a viable bioaugmentation strategy, harnessing the plastic-degrading capabilities of lipase-producing bacteria and fungi. Progress has been made in the field, but questions remain. To date, researchers have not identified *Pseudomonas* spp. lipase degradation of PET ^22,23^, though a cutinase from *P. mendocina* was shown to have high affinity to low crystalline PET, with 5% reduction in the mass of samples ^24^. To our knowledge, our report is the first to find lipase producing *Pseudomonas* isolates that degrades PET, as well as one *Pseudomonas* isolate with the capability to degrade both PET and LDPE plastics. These data might indicate that multiple enzymatic pathways could be involved in degradation. Additionally, the use of *Pseudomonas* spp. are a logical choice for developing bioaugmentation strategies to degrade different types of plastic because of the ability of the genus to respire using difficult to degrade compounds as carbon and energy sources ^25^. While evidence of gross PET and LDPE plastic will take longer to manifest, the initial stages of plastic degradation can be monitored and assessed with the methods presented herein.

Here, two isolates of *Pseudomonas putida*, and a *Pseudomonas chlororaphis* identified by gram staining and 16S ribosomal DNA sequencing, were isolated from the contaminated shoreline of East Galveston Beach, TX. These three isolates were identified through a lipase screen, and Isolate 9.2 appeared to produce more lipase activity than the others (Fig. 3).

However, as identification of lipase activity is not an absolute confirmation of an ability to degrade plastic, all three lipase-positive consortia (9, 10 and 13) were incubated with plastic strips to confirm biofilm formation.

The ability to form biofilms is essential to colonization of plastic ^26^. Biofilms allow for the attachment and protection of bacteria in a scaffold-like matrix and are primarily made up of exopolymeric substance (EPS) composed of polysaccharides, lipids, nucleic acids, and proteins^29^. For instance, biofilm formation on medical devices such as catheters, hip joints, and prosthetic heart valves has been widely reported ^27,28^. Polymers such as glycopeptides, lipopolysaccharides and lipids act as a scaffold that links the biofilm together ^30^.

Evidence of pili via SEM, in conjunction with the sequencing data identifying the isolates as *Pseudomonas putida*, suggest the Pseudomonads in Consortium 9 and Isolate 10 were most likely using the previously characterized Type IV pili (TFP) system for biofilm development and colonization of the PET plastic. Type IV pili are the only pili common to *Pseudomonas*, and in fact most Gram-negative bacteria have them ^31^. TFP are spindly, fibrous organelles found on the surface of many gram-negative bacteria. They are typically involved in bacterial movement on solid surfaces through a twitching motility, as well as bacterial attachment to host cells and extracellular or environmental surfaces ^32^. Additionally, TFP have been shown to be involved in the uptake of macromolecules, as demonstrated by its role in transforming DNA into *N. gonorrheoeae* bacterial cells ^33^. For many bacteria, TFPs are an essential component for biofilm formation, as evidenced by TFP knockout *Psuedomonas aeruginosa’s* failure to build up multicell layers of biofilm on a solid surface ^34,35^. Here, the TFP are associated with islands of EPS as observed with SEM, and bacteria can be found embedded in these rudimentary biofilms (Fig. 6B).

Pili were not observed in SEM imaging of Consortium 13, though it contained *Pseudomonas putida* by 16S sequencing. This could be explained by particular mutations in this isolate or because not all *Pseudomonads* have a TFP system. For example, some *Pseudomonas putida* lack all the subunits necessary to make functional pili and their surfaces lack these structures ^36^. This lack of functional TFP could explain why Consortium 13, which contained one lipase producer and one Gram-positive rod, struggled to colonize PET. The number of bacteria adherent to the surface of the PET was minimal and the EPS production was less than that of the other isolates, observed by SEM (Fig. 6). In screening for plastic-degrading bacteria, this further reinforces the need to test for lipase production and the physical ability to colonize the plastic.

Aiding colonization through the addition of biosurfactants could assist in creating biofilms. Biosurfactants have been shown to both promote and antagonize biofilm formation by allowing for initial colony formation and maintaining nutrient channels essential for a productive mature biofilm, and then promoting their dissolution once migration is necessary ^37,38^. Biosurfactants have the added benefit of increasing hydrophobic surface area to not only aid in the attachment of bacteria, but also to enhance polymer solubility throughout the degradation process ^10,39^ Synthetic biosurfactants like mineral oil can also aid in the colonization and degradation of plastic^40^. Though we have not tested for its presence, it is possible that the *Bacillus cereus* isolates identified as members of Consortia 9 and 13 produce biosurfactants, aiding in colonization and PET degradation.

While these experiments mainly focused on single isolates or consortia with single lipase positive strains, viable bioaugmentation strategies will most likely require the concerted effort of a master consortium of bacteria. Given the many types of plastics to degrade, and the possibility of forming complex biofilms that can degrade plastic faster than any single species, well formulated consortia are the most viable bioaugmentation strategy. It has already been shown that a naturally selected plastic-degrading consortium can degrade plastic faster than individual isolates of that consortium ^41^. Consistently, we found that a consortium of *B. cereus* and *P. chororaphis* exhibited greater lipase activity than singly cultured *Pseudomonas* isolates (Table 2). Further, consortium-based approaches to bioaugmentation pollution clean-up have been successfully utilized before, such as in the aftermath of the Deepwater Horizon oil spill ^42^.

Pretreatment of plastic with UV radiation has been shown to enhance biodegradation of some plastics through free radical formation and introduction of ester functionality into the hydrocarbon backbone of plastic polymers ^7^. This work further supports those results as evidenced by the values in Table 1 where substantial chemical changes were only observed for lipase-positive treatments of PET strips pretreated with UV prior to inoculation. While plastic samples in this study were exposed for 30 minutes, some biodegradation studies have exposed plastic polymers to UV light for up to 8 weeks at 365 nm ^43,44^.

Further work is also underway with increased incubation times to observe larger spectroscopic and gravimetric changes, and compare with other plastic degradation studies carried out over 6 months or longer ^45,46^. When compared to either blank or *E. coli* treatments, PET incubation with all three lipase positive bacteria showed 1) a significant decrease in both the carbonyl and ester indexes with 2) an increase in the aliphatic index. Consortium 13 showed the most promising results for PET degradation (Table 1, Fig. 5B), but also the largest absolute error which may explained by incomplete biofilm formation and subsequent colonization.

We also incubated the lipase positive bacteria with HDPE and LDPE from milk jugs and plastic bags, respectively. Significant changes were not observed in any lipase positive HDPE samples. However, Isolate 10 showed similarly promising results in degrading LDPE and PET (Fig. S1 and Table S2). We focused on the degradation of PET, but these data indicated that the isolates varied not only in their PET degrading capabilities, but also the capability to degrade different types of plastic pollutants. The lipase screen in this work used olive oil as the carbon-source, which is primarily composed of oleic acid, a monounsaturated fatty acid (C18:1). Thus, “lipase-positive” isolates were selected for the ability to utilize primarily ester (-COOR) or midchain olefin (-CH=CH-) groups as a food source. Given the chemical properties of olive-oil are more similar to PET than LDPE or HDPE, it is not surprising the lipase-positive bacteria identified in this work were shown to have lower activity with PE. Furthermore, PE derivatives are much more hydrophobic than PET due to a lack of abundant C-O bonds, which could inhibit biofilm formation and adhesion required for optimal active-site binding from the secreted enzymes. Moreover, biodegradation pathways for PE are not predicted to be the same as for PET and may require a monooxygenase as opposed to a lipase to induce significant biodegradation ^20^.

The degradation pathway for PET is not as well studied as the pathway for PE degradation, but some mechanistic studies have identified how certain bacteria and fungi are able to utilize PET as a sole carbon source ^6^. In general, a major product of PET degradation is the monomer mono(2-hydroxyethyl) terephthalic acid (MHET) and eventually bis(2-hydroxyethyl) terephthalic acid (BHET). PETase is secreted and breaks down PET to MHET, which is broken down by secreted MHETase to terephthalic acid (TPA) and ethylene glycol. Future studies will be necessary to identify the byproducts produced during PET degradation by the Pseudomonads isolated in this study.

Ideally, a viable consortium-based bioaugmentation strategy would consist of growing plastic degrading bacteria within a contained, carbon-free system. This would ensure that bacteria would utilize and degrade only plastic waste that was introduced into the system. Pre-treatment of plastic would occur prior to bacterial degradation to render the inert polymer more amenable to bacterial degradation. Specifically, plastic waste would be subjected to UV-pretreatments, or other means to introduce ester linkages into the inert polymer backbone that are more easily recognized and cleavable by bacterial lipases. The waste would also be subjected to mechanochemical grinding into smaller fragments, resulting in more surface area for the bacteria to colonize. The plastic waste would then be fed into the contained system to be degraded. End products from the process would include bacterial biomass, that could be used as fertilizer, and carbon dioxide. While much work is needed to bring bacterial degradation of waste plastics to the industrial scale, our work, and that of others indicate that bioaugmentation is a viable strategy to help mitigate the immense pollution problem that we humans have created.

## MATERIALS AND METHODS

### Soil sample collection

Superfund sites are sites that are so polluted the EPA has made their clean-up a national priority; eleven superfund sites exist in the greater Houston, TX area. Using the logic that bacteria in polluted environments are more likely to adapt to harnessing pollutants to survive, soil samples were collected from eight different sites along the gulf coast of southeast Texas and within the greater Houston area. Soil samples (500g) were collected from each of the following sites. Sample 1 was collected at the Jones Road Chemical Plume Superfund Site (Houston, TX), currently home to a large strip-mall and apartment complex despite high levels of petroleum-based dry-cleaning solvents in the soil ^47^. Sample 2 was collected at the Baer Road Foundry Superfund Site (Houston, TX), home to a steel manufacturing plant for 70 years where high levels of heavy metal deposits and petroleum byproducts have been found in 34% of nearby homes ^47^. Samples 3, 4 and 5 were collected at the Pasadena Refining System, a large petrochemical industrial complex along the Houston Ship Channel that processes roughly 106,000 barrels of crude oil per day ^48^. Samples 6 and 8 were collected at the West Park and Baer Road Power Stations (Houston, TX), respectively, on a network of buried wires coated in petroleum-based insulation coating ^49^. Sample 7 was collected near the shoreline at East Beach (Galveston, TX) and provided the richest source of plastic degrading bacteria. Sample 8 was collected near the transformers of the West Park Power Station (Houston, TX). All samples were collected roughly 6 inches beneath the topsoil layer and refrigerated prior to transport in sealed zip-lock bags.

### Bacterial extraction from soil

Each soil sample (2g) was resuspended in 9mL Phosphate buffered saline (PBS) prepared accordingly per 1-liter diH_2_O: 8g NaCl, 0.2g KCl, 1.44g Na_2_HPO_4_, and 0.24g KH_2_PO_4_ adjusted to pH 7.4 and autoclaved for 20 minutes (15 psi, 121°C). The soil and PBS suspensions were placed on rotary shaker (250rpm) for 24 hours. The sediment was allowed to settle, and 100μL of this suspension was then spread on Lysogeny Broth (LB) agar plates prepared accordingly per 1-liter diH_2_O: 10 g Tryptone, 5g Yeast, 5g NaCl, 18 g agar adjusted to pH 7 and autoclaved (15 psi, 121° C). Multiple plates were made from each soil, inverted and incubated at 26° C for 24 hours.

### Lipase screening

Rhodamine B agar plates (per 1 liter: 9.0 g nutrient broth powder, 2.5g yeast extract and 10g agar) were prepared to test isolated bacterial colonies for lipase activity. For the lipid emulsion media, 250 μl of Tween80 and 30ml olive oil were added to 50ml diH_2_O and emulsified in a blender. The final lipoidal emulsion was adjusted to pH 7. The base media and lipoidal emulsion were autoclaved separately. Following autoclaving, Rhodamine B was added to a concentration of 0.024% (w/v) to the sterile lipoidal emulsion. Lipoidal emulsion (50 ml) was then added to the base nutrient media to a final volume of 1 liter, mixed thoroughly, with the final concentration of Rhodamine B at 0.0012% (w/v), and then poured^50^.

Colonies grown on LB agar were screened for bacterial lipolytic activity via a colony lift assay from the LB plate to Rhodamine B agar. The Rhodamine B plates were inverted and incubated for 24 hours at 26 °C. Colonies producing lipase on Rhodamine B agar were identified as those that produced fluorescent halos when exposed to a UV trans-illuminator at 365nm and were re-streaked onto individual LB plates for isolation and purification (see Figure 1). *E. coli* MC4100 was used as a negative control. This assay was repeated to ensure isolated strains remained lipase positive.

### Purifying cultures

One hundred ninety-two colonies were screened for lipase activity on Rhodamine B plates. One hundred seventy-eight were negative, with 14 colonies initially being positive. Three colonies remained lipase positive on Rhodamine B plates. Overnight cultures of the 3 bacterial colonies were repeatedly streaked on LB agar to obtain pure cultures. Gram staining was utilized to help confirm bacterial strain purity and to corroborate 16S PCR results.

### Gram Staining

One sterile loop of liquid culture (OD_600_ = 1.0) was spread onto sterile glass slides and heat fixed. The slide was flooded with crystal violet for 1 minute, washed with diH_2_O for 5 seconds, flooded with Gram’s iodine for 1 minute, washed with diH_2_O for 5 seconds, flooded with 95% EtOH for 10 seconds and flooded with safranin for 1 minute, prior to a final diH_2_O rinse and blotting with bibulous paper. Slides were visualized using 1000x magnification. Images were captured using a Keyence BZ-X700 inverted fluorescence and color microscope.

### 16S rRNA gene PCR and DNA sequencing

Colony PCR was performed using universal 16S primers (Table S3) on the three lipase-positive strains and two additional isolates from Consortia 9 and 13, using a standard thermocycler program ^51^. The PCR products were imaged in a 1.2% agarose gel in TAE buffer run at 110 mV for 30 minutes. PCR products were cleaned using a GENECLEAN (MP Biomedicals, Santa Ana, CA) and resuspended in sterile water.

Following DNA sequencing at ACGT, INC. (http://www.acgtinc.com/) sequences were aligned using BioEdit 7.2.5 biological sequence alignment editor. Prior to alignment, chromatograms were checked to ensure quality sequences and the first ~25-30 nucleotides from each sequence were eliminated due to sequencing artifact. Gaps were inserted manually until maximum alignment had been achieved. No chimeric sequences were observed. Only the core sequence with 100% agreement (150-764 nucleotides) was used to determine genus identity. Genus identification was done using nucleotide BLAST (BLASTn), and identity cutoffs were set to only those matching 100%. The 100% identity metric was employed due to the conserved nature of the 16S rRNA gene targeted by these primers and conserved regions of the 16S rRNA gene can tolerate very few base pair changes and thus a single base pair may be the difference between two genera^51^. Primers directed to the V3-V6 region of the 16S rRNA gene were used for identification at the species level^52^.

### Random Amplification of Polymorphic DNA (RAPD) PCR Analysis

RAPD PCR was conducted on each of the five bacterial isolates using the ERIC2 primer (5’-AAGTAAGTGACTGGGGTGAGCG-3’, 57°C annealing)^17^. Products were imaged in a 1.8% agarose gel.

### Liquid carbon free media (LCFM) supplemented with plastic

Carbon free base media was prepared accordingly per 1 L of diH_2_O: 0.7 g KH_2_PO_4_, 0.7 g K_2_HPO_4_, 1.0 g NH_4_NO_3_, and 0.005 g NaCl, to ensure plastic strips were the sole source of carbon available during the incubation. A 1 M stock solution of essential metals was prepared accordingly in 100 ml of sterile H_2_O: 7 g MgSO_4_·7H_2_O, 20 mg Fe_2_SO7.H_2_O, 20 mg ZnSO_4_·7H_2_O, and 9 mg MnSO_4_·H_2_O (stirred 4 h), after which 10 ml were filter sterilized and added to 1 liter autoclaved liquid base media. Polyethylene terephthalate (PET), high-density polyethylene (HDPE), and low-density polyethylene (LDPE) samples were cut into 25 x 5 mm strips, sterilized in 70% EtOH and hung in a Biosafety cabinet to dry.

Cultures of the three lipase positive consortia were grown overnight in LB and diluted to an OD_600_ of 1 to ensure equal amounts of bacteria were added to each sample. For single point LCFM incubations, 10 uL of overnight culture was added to each 4 ml tube of LCFM.

Previously weighed and sterilized plastic strips were added to each tube (1 type per tube). The samples were incubated in an environmental shaker (26 °C, 125 rpm) for 3 months. Samples were replenished with sterile LCFM each month due to evaporation.

Another set of LCFM cultures was set up following the same above procedure with and without UV irradiation of the PET, HDPE, and LDPE strips with a 365 nm UV lamp for 30 minutes prior to sterilization and inoculation. Each test tube was filled with 8 ml LCFM and inoculated with 50 μL overnight culture (OD=600). Tubes were set up in triplicate and for static incubation at 26 °C for six weeks, to increase biofilm formation.

### Surface characterization with attenuated total reflectance – Fourier transform infrared spectroscopy (ATR-FTIR)

Plastic PET strips were submerged in 30 ml 2% SDS and placed on the rotary shaker for 2 hours (225 rpm, 37°C) to remove biofilms, immersed in fresh diH_2_O water and air-dried. A ThermoScientific iS5 infrared spectrometer and id7 diamond-ATR attachment was used to acquire spectra from 4000 – 450 cm^−1^ (4 cm^−1^ resolution) with Omnic software. Data were transformed using an N-B strong apodization and Mertz phase correction. Three areas were analyzed for each sample to obtain spectra that represent the average condition of the plastic surface. All infrared spectra were normalized by peak intensity to common C-H bending modes used for spectral normalization of these polymers: 1409 cm^−1^ for PET and 1368 cm^−1^ for LDPE and HDPE.

### Biofilm imaging by scanning electron microscopy (SEM)

Plastic samples were soaked in 2% phosphate buffered glutaraldehyde for cell fixation. For post-fixation, samples were submerged in 2% osmium tetra-oxide in an ice bath for 3 hours. The samples were then dehydrated in graded EtOH (50, 75 and 100%) baths for 15 minutes each before undergoing critical point drying with CO_2_. Dried samples were sputter-coated with gold using a Leica ACE600 Coater prior to imaging with a FEI Helios Nanolab 660 DualBeam FIB-SEM and TLD detector operating at an accelerating voltage of 2 kV with 1 μs dwell time.

### Statistics

Peak intensity ratios for ATR-FTIR were calculated using the average intensities from nine spectra (3 measurements x 3 samples for each condition) and standard deviations were used to propagate errors prior to calculation of confidence intervals (n=9, α=0.05). Studies performed in biological triplicate were compared by unpaired two-tailed Student’s t-test or 1-way ANOVA with replication and Tukey’s HSD. Values of p<0.05 were considered statistically significantly different.

### Strain Deposition

The following bacterial isolates were deposited with the USDA NRRL

|  |  |  |  |
| --- | --- | --- | --- |
| Culture Collection: | NRRL B-67630 | R10 | *Pseudomonas putida* |
|  | NRRL B-67631 | R131 | *Bacillus cereus* |
|  | NRRL B-67632 | R91 | *Bacillus cereus* |
|  | NRRL B-67633 | R92 | *Pseudomonas chlororaphis* |
|  | NRRL B-67634 | R132 | *Pseudomonas putida* |

## ACKNOWLEDGEMENTS

The authors thank Claudia S. López, PhD, Director of the Multiscale Microscopy Core at Oregon Health & Science University for scanning electron microscopy imaging. This work was, in part, funded by an undergraduate research fellowship awarded to MV by Reed College.

## AUTHOR INFORMATION

### Contributions

J.L.M., M.V., and N.A.S. wrote the manuscript. M.V. isolated and identified lipase-positive and consortia bacteria, established plastic cultures and prepared samples for ATR-FTIR spectroscopy. C.R. aided in bacterial species identification and compared lipase-producing isolates, while G.C. assisted in ATR-FTIR analysis. All authors reviewed the manuscript.

### Competing interests

The authors declare no competing interests.

**Corresponding author**

Correspondence to Jay L. Mellies.

